# Probing Ligand and Cation Binding Sites in G-Quadruplex Nucleic Acids by Mass Spectrometry and Electron Photodetachment Dissociation Sequencing

**DOI:** 10.1101/563627

**Authors:** Dababrata Paul, Adrien Marchand, Daniela Verga, Marie-Paule Teulade-Fichou, Sophie Bombard, Frédéric Rosu, Valérie Gabelica

## Abstract

Mass spectrometry provides exquisite detail on ligand and cation binding stoichiometries with a DNA target. The next important step is to develop reliable methods to determine the cation and ligand binding sites in each complex separated by the mass spectrometer. To circumvent the caveat of ligand derivatization for cross-linking, which may alter the ligand binding mode, we explored a tandem mass spectrometry (MS/MS) method that does not require ligand derivatization, and is therefore also applicable to localize metal cations. By obtaining more negative charge states for the complexes using supercharging agents, and by creating radical ions by electron photodetachment, oligonucleotide bonds become weaker than the DNA-cation or DNA-ligand noncovalent bonds upon collision-induced dissociation of the radicals. This electron photodetachment (EPD) method allows to locate the binding regions of cations and ligands by top-down sequencing of the oligonucleotide target. The very potent G-quadruplex ligands 360A and PhenDC3 were found to replace a potassium cation and bind close to the central loop of 4-repeat human telomeric sequences.

## INTRODUCTION

In the presence of physiologically relevant cations such as potassium, guanine-rich repeated sequences can form G-quadruplex structures owing to the formation of guanine quartets, which pile up thanks to monovalent cation coordination in-between them (Figure 1).^1^ Sequences having at least four G-rich regions can form a variety of intramolecular G-quadruplex topologies, as the three loops linking the G-tracts can adopt no less than 14 different combinations of lateral, diagonal, or edgewise positioning.^2, 3^ This structural polymorphism is what renders G-quadruplex structural studies difficult. The polymorphism is particularly acute for the human telomeric motif, constituted by repetitive (TTAGGG)_n_ sequences, because the 3-nt loops (TTA) are compatible with edgewise, lateral and diagonal position, and thus multiple topologies are possible.^4–8^ Moreover, these conformations are relatively close in energy, and minor changes in the sequence (e.g., addition or deletion of terminal bases^4, 7^) or in the solution conditions (e.g., the nature monovalent cations present,^8^ or the presence of non-aqueous additives^9, 10^) can switch the conformational equilibria from one topology to another. Finally, structures with two G-quartets can be formed alongside those having three G-quartets.^11, 12^

**Figure 1.**
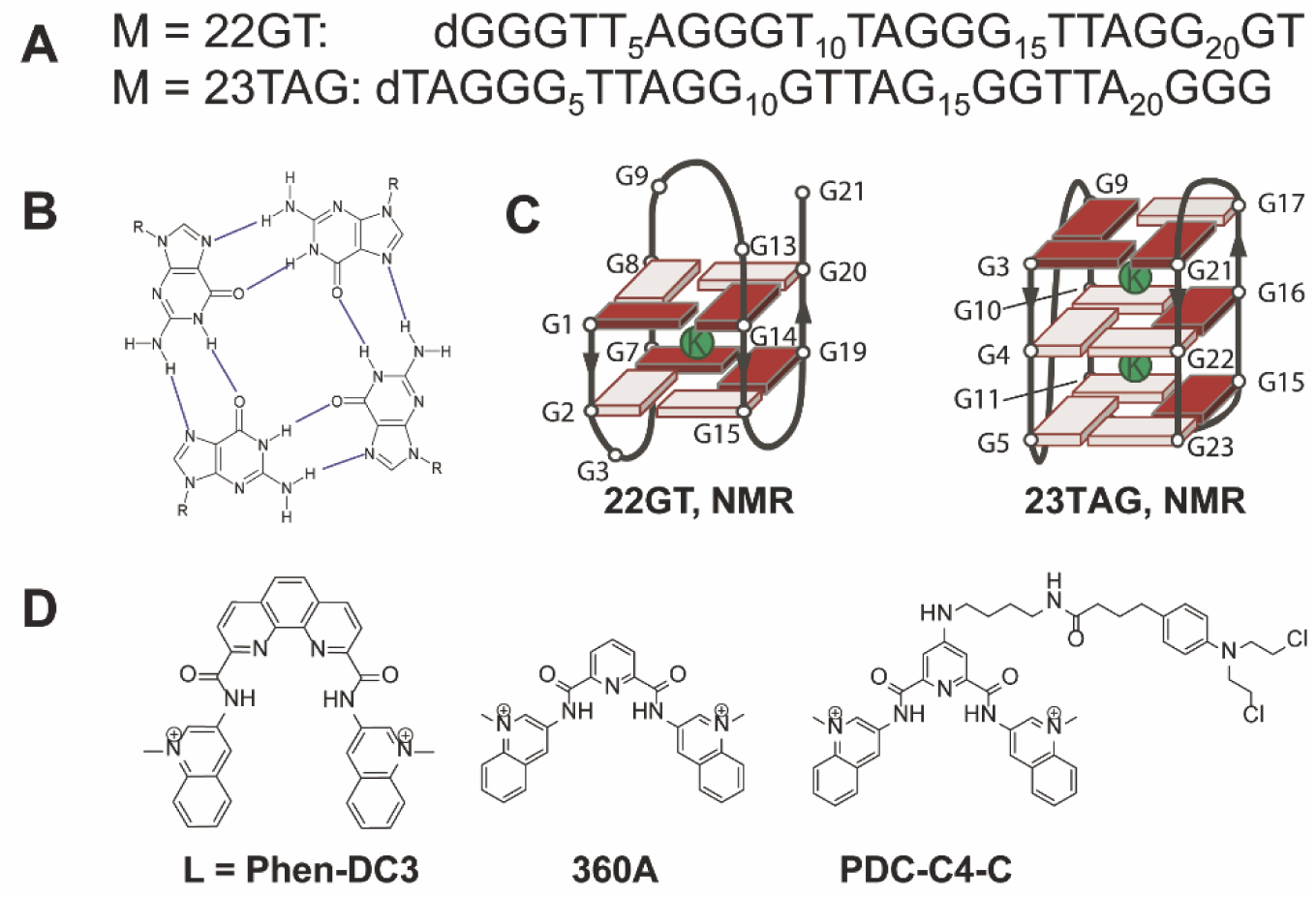
A) G-quadruplex DNA forming sequences studied here. B) Hydrogen bonding pattern in a guanine quartet. C) Schematic topology adopted by the sequences 22GT and 23TAG, as reported from solution NMR (PDB accession codes: for 22GT:2KF8^11^, for 23TAG: 2JSM^24^), and supposed potassium ion locations, sandwiched in-between the G-quartets. White grey rectangles are guanines in *anti* conformation, and red rectangles are guanines in *syn* conformation. D) Chemical structures of the ligands studied herein.

G-quadruplex formation can influence important cellular processes such as gene expression (particularly for oncogenes),^13^ splicing,^14^ or telomere maintenance.^15, 16^ The search for ligands able to bind specifically to particular G-quadruplexes is therefore a very active field of research, either for therapeutic aims^17–19^ or to probe G-quadruplex presence.^20^ The rational design of drugs targeting G-quadruplexes is also hampered by the fact that organic molecules can also shift conformational equilibria. In the context of G-quadruplex ligand design, it is important to characterize experimentally not only ligand binding affinities, but also ligand binding modes including ligand-induced conformational changes.

We have previously demonstrated using mass spectrometry (MS) and circular dichroism (CD) spectroscopy that the high-affinity G-quadruplex ligands PhenDC3 and 360A (Figure 1D) change the conformation of the human telomeric G-quadruplexes by ejecting a cation upon binding: the major DNA(M):ligand(L):cation(K) stoichiometry is 1:1:1 (the complex is noted “MLK”), despite in the same conditions without ligand the main DNA:cation stoichiometry is 1:2 (complex “MK_2_”).^21–23^ However, the ligand binding site, (i.e. the G-quartet), cannot be determined by simple MS experiments. Here, we used mass spectrometry to select complexes of well-defined cation and ligand binding stoichiometries, and explored a top-down fragmentation approach (i.e. fragmentation of the intact complex) to locate the non-covalently bound cations and ligands on the target sequences.

To reliably locate a noncovalently bound ligand on a DNA sequence using tandem mass spectrometry (MS/MS), ligand-DNA bonds must not break while the DNA backbone must fragment at multiple positions to obtain full sequence coverage. This objective is identical as for the localization of fragile post-translational modifications on proteins^25^ Two approaches (which can be combined) maximize the chances of success: (1) reinforcing the bonds between the noncovalent ligand and the DNA, and (2) weakening the DNA backbone to favor chain fragmentation.

In electrospray ionization (ESI) mass spectrometry, nucleic acids spray well in the negative ion mode, and most high-affinity DNA ligands are positively charged, so ligand-DNA ionic interactions are actually reinforced in the desolvated ions compared to in solution, and may persist even under the CID (collision-induced dissociation) conditions required to fragment the DNA backbone. This property has been exploited to characterize aminoglycoside binding to RNA^26^ and tat peptide binding to TAR RNA^27^ by CID. Another way to reinforce bonds is to derivatize the ligand with an alkylating agent that will irreversibly bind to DNA bases accessible from the binding site. To accurately reflect the ligand binding properties, the derivatization must not perturb the binding site and the conformational equilibria. The alkylated base(s) can then be mapped by gel electrophoresis, or by MS/MS. The widespread collision-induced dissociation MS/MS technique typically leads to fragmentation into *w* and *a-Base* ions (Figure 2A,B).^28^

**Figure 2.**
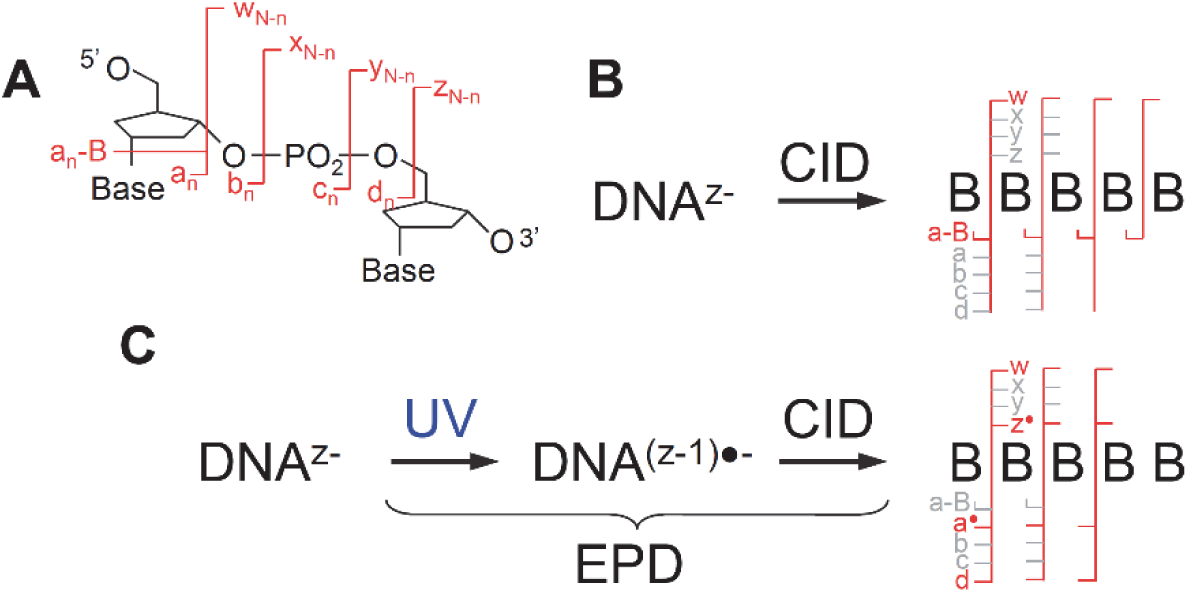
A) Fragment ion nomenclature for oligonucleotides. B) Typical fragments encountered in collision-induced dissociation (CID) of closed-shell DNA: *a_n_-B* (with loss of base *n* from fragment *a*_*n*_) and *w_N-n_*, and schematic annotations for CID sequencing (*n* indicates fragmentation after the n^th^ base, and *N* is the total number of bases in the oligonucleotide). C) The electron photodetachment dissociation (EPD) process (CID on radicals created by UV irradiation), typical fragments (*a*_*n*_^●^, *w_N-n_*, *d*_*n*_ and *z_N-n_^●^*) observed in EPD of oligonucleotides, and schematic annotations for EPD sequencing.

The second approach is to use alternative MS/MS activation techniques that weaken the DNA backbone. Typically, radical ions are less stable than their corresponding closed-shell ions, and sequence coverage is more extensive from radicals. In proteins, creating radicals by electron capture dissociation (ECD) or electron transfer dissociation (ETD) are widely used in top-down sequencing of multiply charged cations, and these approaches have been used to locate cation^29^ or ligand binding sites.^30–33^ Nucleic acids, however, are best ionized as multiply charged anions. Radicals are therefore more easily accessed by electron detachment.^34^ Irradiation of DNA multiply charged anions with UV light (235-280 nm) is particularly efficient to detach electrons.^35^ A recent systematic study on 6-mer single strands shows that the competition between electron photodetachment (ePD) and fragmentation (UVMPD) is base-dependent, and that electron photodetachment is particularly efficient for guanines.^36^ However, for longer strands, UVMPD becomes inefficient, most probably because of the competing collisional cooling by the helium in the quadrupole ion trap, and further activation is required to obtain fragments. The radicals generated can thus be selected in an MS^3^ step and fragmented by CID. The whole process, illustrated in Figure 2C, has been coined electron photodetachment dissociation (EPD).^37^

EPD sequencing has been demonstrated first for DNA, with complete coverage for single strands up to 15 nucleotides,^37^ and then has been applied to proteins,^38^ carbohydrates, and synthetic polymers.^39^ This method is appealing for characterizing non-covalent interactions between ligands and G-quadruplexes because (i) G-quadruplexes ionize well in negative mode and the presence of several guanines ensures efficient photodetachment, and (ii) electron photodetachment is soft and the structure of the complexes is likely preserved during that step. The challenge here is to obtain as complete sequence coverage as possible on sequences longer than 20 nucleotides, and to validate that the cations and ligands do not shuffle before the fragments separate at the MS^3^ step. Here, we explored whether EPD could provide information on the cation and ligand binding sites on the G-quadruplexes, with particular focus on characterizing the peculiar binding mode of PhenDC3 and 360A to telomeric G-quadruplexes. We found that these ligands bind close to the central loop of the quadruplexes, and validated this result with a covalent derivative.

## MATERIALS AND METHODS

### Samples

All oligonucleotides were purchased from Eurogentec (Seraing, Belgium) in lyophilized form and with RP-cartridge-Gold purification. They were solubilized in nuclease-free grade water (Ambion, Fisher Scientific, Illkirch, France) at approximately 200 µM. The concentrations were measured using absorbance at 260 nm on a Uvikon XS and molar extinction coefficients calculated using the online Integrated DNA Technologies OligoAnalyzer tool using Cavaluzzi-Borer Correction.^40^ Final solutions at the exact desired concentrations were prepared in nuclease-free grade water with trimethylammonium acetate (TMAA, Ultra for UPLC, Fluka) and potassium chloride (KCl, >99.999%, Sigma) purchased from Sigma-Aldrich (Saint-Quentin Fallavier, France). The experiments were carried out on the sequences 22GT and 23TAG (Figure 1A), which predominantly adopt specific topologies in KCl solution (Figure 1C). These sequences were prepared in 0.2 mM to 1.0 mM KCl and 100 mM trimethylammonium acetate (TMAA) solutions, which mimic physiological ionic strength, ensure K^+^ presence to fold the G-quadruplexes, and are compatible with electrospray ionization mass spectrometry.^41^ The synthesis of the covalent binder analogues is described in the supplementary information.

### Mass spectrometry and electron photodetachment dissociation (EPD)

ESI-MS, MS/MS and MS^3^ experiments were carried out on a Bruker Amazon SL quadrupole ion trap mass spectrometer (Bruker Daltonics, Bremen, Germany) modified to allow a laser beam to make a single pass through the center of the trap. The electrospray source conditions were 3000 V on the capillary, 500 V on the end plate offset, 7.2 psi on the nebulizer, and 3.5 L/min and 180 °C on the dry gas and dry temperature, respectively. Each mass spectrum was recorded for 10 minutes, and over 40 minutes were averaged to construct each EPD spectrum. A PREMISCAN/500/MB OPO with UV scan (GWU, Erftstadt, Germany), pumped by a Quanta-Ray PRO-230-30 Nd:YAG laser (Spectra Physics, Santa Clara, USA), was used to produce ~2 mJ/pulse at 240 nm and 30 Hz. This wavelength was chosen to maximize electron photodetachment, but note that any wavelength in the base absorption region (~235-280 nm) would produce EPD as well. The EPD experiments were carried out by synchronizing a mechanical shutter opening with the activation time (100 ms, i.e. 3 laser shots) of the MS^2^ event. The laser beam (typical energy: ~1mJ/pulse at the vacuum chamber window) was focused on the ion cloud by a lens of 700 mm focal length. Three laser pulses per duty cycle were used to produce radical anions at 0V CID activation amplitude. The resulting radical anions were mass-isolated (MS^3^ stage, 4 *m/z* window) and subjected to collision-induced dissociation (typically: 1.2 V). The instrument was operated in negative ion mode (capillary voltage 3000 V, capillary exit −140 V). The syringe pump flow rate was 180 µL/h. EPD spectra were extracted using Data Analysis (Bruker Daltonics) software and interpreted manually.

### Ion mobility spectrometry

Ion mobility (ESI-IM-MS) experiments were carried out on an Agilent 6560 IMS-Q-TOF (Agilent Technologies, Santa Clara, USA) operated with helium in the linear drift tube IM. The dual-ESI source was operated in the negative ion mode. The syringe pump flow rate was 180 µL/h. The instrument is equipped with the “Alternate Gas Option”, wherein capacitance diaphragm gauges are connected to the trapping funnel and to the drift tube, and an additional flow controller admits gas in the trapping funnel, and the flow controller is regulated by a feedback reading of the pressure in the drift tube. An in-house modification to the pumping system allows faster equilibration of the pressures: a second Tri-scroll 800 pump (Agilent) was connected to the source region (with an Edwards SP16K connected to the front pumping line), while the original Tri-scroll 800 pump is connected to the Q-TOF region. For all measurements, the helium pressure in the drift tube was 3.89 ± 0.01 Torr, and the pressure in the trapping funnel is 3.67 ± 0.01 Torr. The source temperature and fragmentor voltage were set at 200°C and 350V. The RF of the high pressure funnel and trapping funnel were set to 200 V and 210 V respectively. To ensure soft trapping condition, the trap entrance grid delta was set to 6 V.

## RESULTS AND DISCUSSION

A typical ESI-MS spectrum obtained for 10 µM 23TAG in 0.2 mM KCl/100 mM TMAA is shown in Figure 3A: at this KCl concentration, the free DNA (M) and the complexes with one or two potassium ions (MK and MK_2_) coexist in approximately equal proportions. This KCl concentration was chosen so that, according to the equilibrium constants of potassium binding to the 23TAG sequence,^42^ both the unfolded form (M) and the two folded forms (MK and MK_2_) are visible, and nonspecific adducts are minimal. In 1 mM KCl, the complex MK_2_ becomes predominant (supporting Figure S1). The main charge states are 4- and 5-. When 10 µM ligand PhenDC3 is added to the 1 mM KCl solution of 23TAG, the major complex formed is however MKL (Figure 3B): the ligand displaces one potassium ion. This ligand-induced K^+^ replacement is observed for all sequences and ligands studied here (supporting Figure S2).

**Figure 3.**
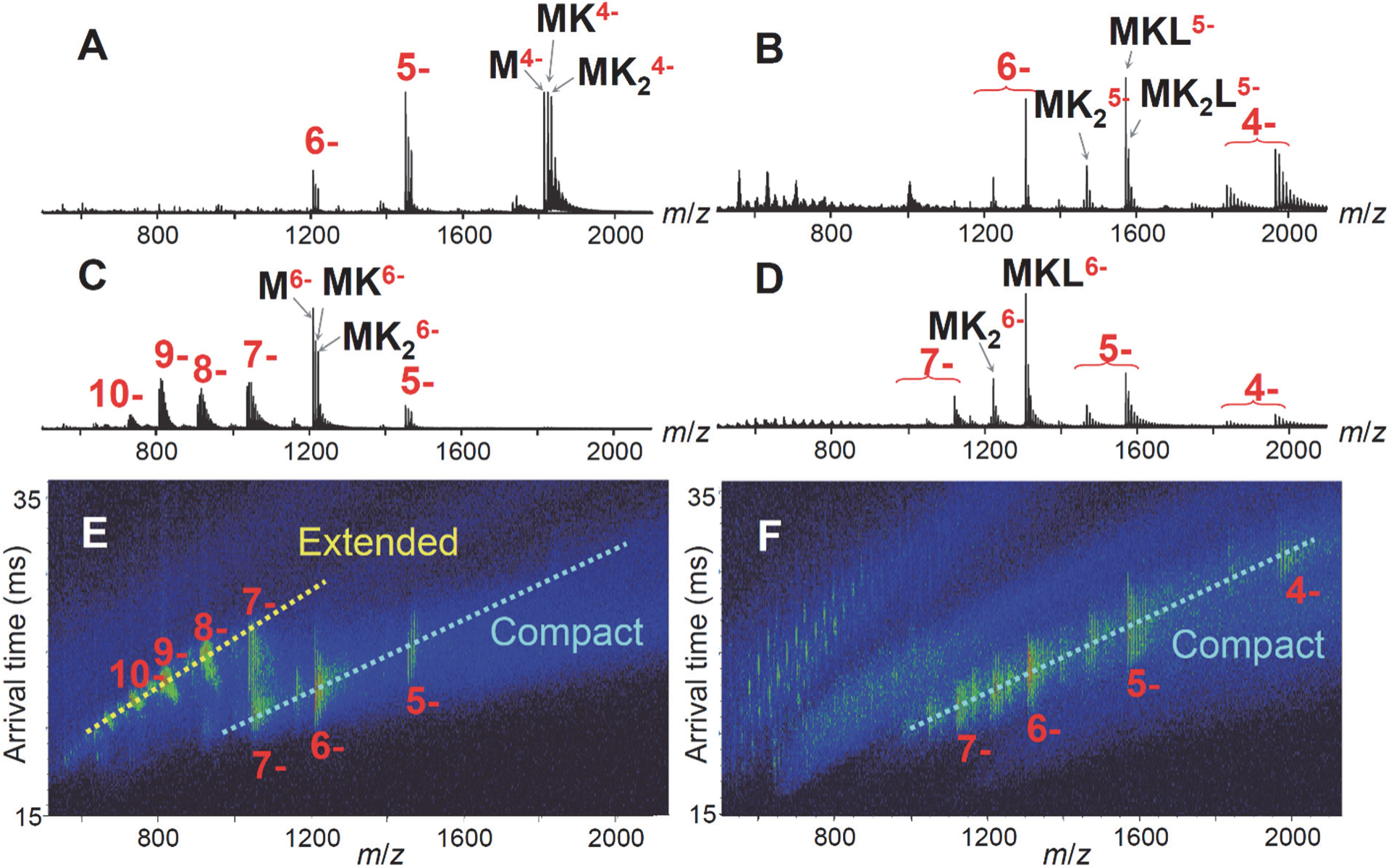
A) ESI-MS spectrum of 10 µM 23TAG in 0.2 mM KCl/100 mM TMAA. B) ESI-MS spectrum of 10 µM 23TAG + 10 µM PhenDC3 in 1.0 mM KCl/100 mM TMAA. C-D) ESI-MS spectra of the same solutions as A-B, with 0.75% (vol) sulfolane added. E-F) Bi-dimensional separation (mass spectrometry on x-axis and ion mobility drift time separation on y-axis) of spectra C-D. The arrival time following drift tube separation is proportional to CCS/z (CCS = collision cross section). Charge state distributions having the same CCS are therefore found on diagonals. Ions of the most compact conformations travel faster.

CID and EPD experiments were all carried out on a quadrupole ion trap instrument modified to couple and synchronize a wavelength-tunable laser operated at 240 nm. All EPD experiments starting with the 5- charge state of the 22-mers resulted in poor sequence coverage (supporting Figure S3): the Coulomb repulsion in the M^4-●^ radical produced at the MS^2^ step is not sufficient to separate the fragments in the MS^3^ step. EPD experiments must be performed on higher charge to start with, but the 6- charge state produced from 100 mM TMAA (or the more typically used 100 mM NH_4_OAc) solutions of the 22/23-mers is not abundant enough to produce EPD spectra with sufficient signal-to-noise ratio.

To increase the abundance of higher charge states, we used “supercharging” additives such as meta-nitrobenzylalcohol (m-NBA) or sulfolane, which increase the net charge of protein cations and nucleic acid anions.^43–47^ All EPD results were obtained with addition of 0.75% volume sulfolane. Importantly, we were concerned that supercharging could also disrupt the complex structure and affect the ligand position before probing it by EPD. Therefore, we first verified by electronic circular dichroism spectroscopy that the sulfolane additive did not affect the folding of the complexes in solution (supporting Figure S4). We found no CD signal change, even at up to 7.5% sulfolane in solution (an extreme situation that could occur upon droplet desolvation, due to the preferential evaporation of water), suggesting that sulfolane did not affect the G-quadruplex topology in solution.

We used ion mobility spectrometry on the solutions prepared with 0.75% m-NBA or sulfolane to verify which charge states remain folded in the gas-phase prior to EPD, and which charge states were too high and caused unfolding already upon electrospray. The ion mobility analysis on sulfolane-doped solutions of 10 µM 23TAG + 0.2 mM KCl (ESI-MS spectrum in Figure 3C) and 23TAG + 1 mM KCl + 10 µM ligand (Figure 3D) are shown in Figures 3E and 3F, respectively. The ion mobility results deserve a few comments. First, both the G-quadruplex folded forms (MK and MK_2_) and the nonfolded form (M) adopt compact gas-phase conformations when produced at charge states 4- to 6-. Actually, for charge state 5-, the nonfolded form is more compact in the gas phase than the G-quadruplex forms. At charge state 7-, the nonfolded form M is mostly extended in the gas phase, while a fraction of the G-quadruplexes (especially MK_2_) remains partially compact. We interpret this behavior as resulting from the balance between Coulomb repulsion and intramolecular noncovalent bonds. At charge state 5-, the Coulomb repulsion is not large enough to compensate for the nonspecific hydrogen bonds that can form between the polar and charged groups of single strands, let alone for the specific G-quadruplex bonds. The gas-phase compaction at low charge states was discussed before for DNA single strands,^48^ DNA and RNA duplexes,^49^ and DNA i-motifs.^50^ At charge state 7-, the nonspecific interactions of the nonfolded forms are mostly broken, whereas those of the G-quadruplex persist up to higher internal energies. Charge states 7- and higher can be obtained only with the addition of sulfolane. Given that sulfolane does not significantly denature the complexes in solution, we suppose that charging occurs first, and then upon removal of the last solvent molecules the dielectric constant decreases and Coulomb repulsion unfolds the strands in the gas phase.

Similar ion mobility results were obtained for all sequences and ligands. Interestingly, the complexes at the charge state 6- remain compact in the gas phase (same collision cross section as the 5- charge state) in all cases. To take no risk, we used only ions of the 6- charge state for probing by EPD, and therefore analyzed the fragments of the 5-● ions.

EPD was carried out first for the complexes with potassium ions. The EPD results for 22GT without potassium (unfolded) show complete sequence coverage (Figure 4A), in contrast with regular CID (Supporting Figure S5), where cleavages on the 3’ side of thymines are often missed.^28^ However, when one potassium ion is bound to 22GT, a portion of the sequence around the central loop (from G9 to G14) is protected from fragmentation (Figure 4C). This also corresponds to the exclusive zone where the potassium ion remains attached upon fragmentation: K^+^ is not detected on 5’-terminal fragments equal to or shorter than *a*_8_^•z-^, and always detected on fragments equal to or longer than *a*_14_^•z-^. Complementarily, potassium is not detected on 3’-terminal fragments up to *w*_8_^z-^ and are detected starting at *w*_15_^z-^.

**Figure 4.**
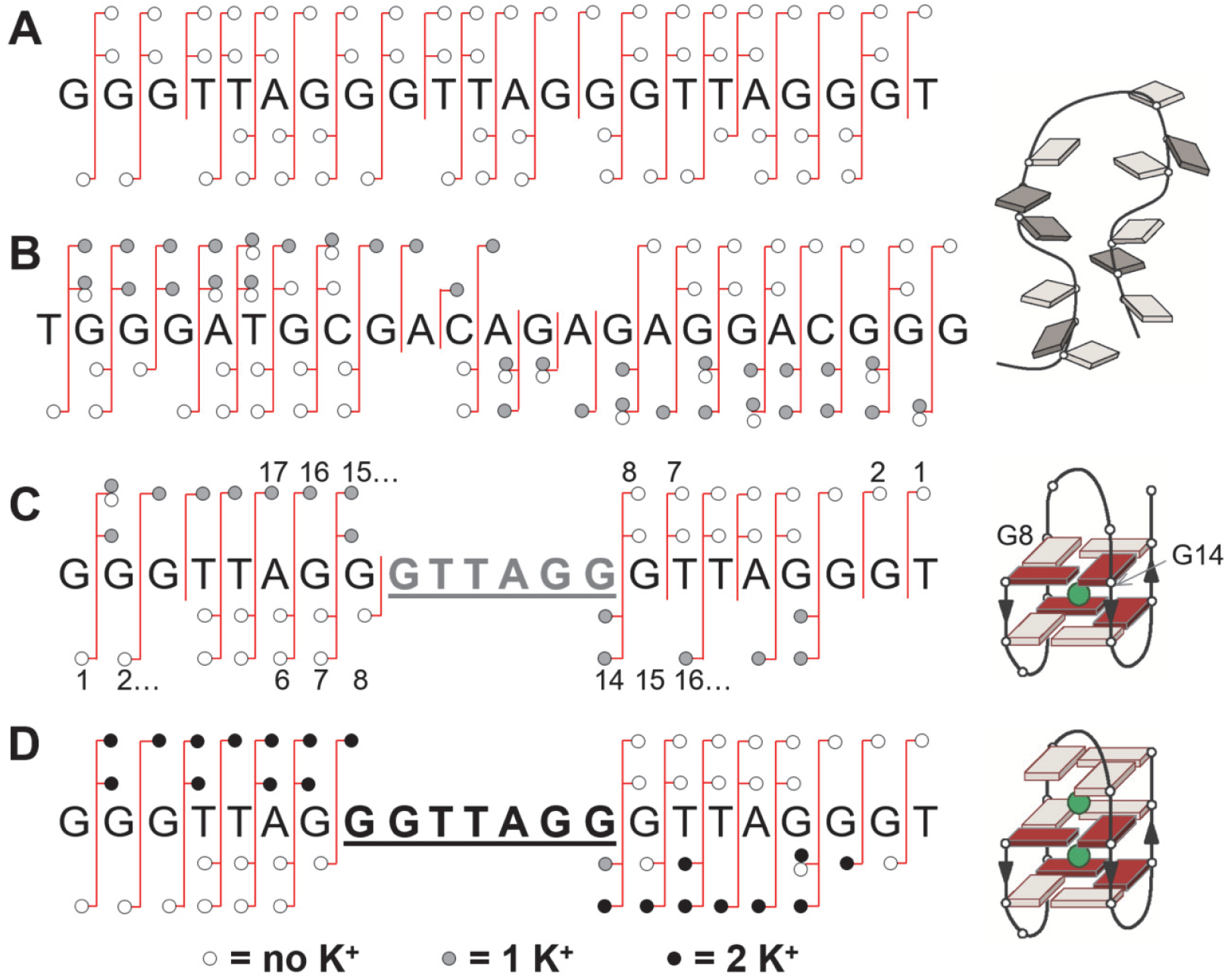
A) EPD analysis of 22GT without potassium ion bound (M^6-^). B) EPD of the nonspecific MK^6-^ complex of a control strand not forming G-quadruplex. EPD of 22GT G-quadruplex with one (C) potassium (MK^6-^) or with (D) two potassium ions bound (MK_2_^6-^). Fragment schematic nomenclature is defined in Figure 2; and key numbering in panel C.

Importantly, the fragmentation of the potassium complex of a control sequence not able to form G-quadruplexes (the potassium adduct has therefore no specific location), gives (1) no such protection and (2) a more random positioning of the K^+^ on the fragments (Figure 4B: a high proportion of fragments are detected both with and without K^+^ bound). The protection and particular K^+^ location is therefore a tenet of the quadruplex fold.

The 22GT sequence is known to fold into a 2-quartet structure formed by G1-G14-G20-G8 and G2-G15-G19-G7. The potassium ion is octa-coordinated between these eight guanine residues. The results therefore suggest that either potassium precludes radical-induced fragmentation at the backbone close to the loop region (but there is no clear rationale for that), or that the fragments formed after backbone cleavage around the central loop cannot separate in the CID activation conditions wherein the other bonds easily break. This is plausible because separating the fragments following cleavage at the central loop involves breaking more K^+^-guanine coordination bonds than separating the fragments following cleavage of the other loops (4 bonds versus 2 bonds). For the observed fragments, the potassium ions always stay on the longer fragment, possibly because they remain coordinated to guanine triplets. EPD of the complex of 22GT with two potassium ions shows a similar protection and location (G8 to G14) of both K^+^ ions close to the central loop (Figure 4D). This is in line with the intuitive anticipation that the additional K^+^ binding site resides between the G1-G14-G20-G8 quartet and the G9-G13-G21 triplet. Results for 23TAG were similar (Supporting Figure S6): for example, for the MK_2_ complex, protection and K^+^ location was at the exact same central sequence GGTTAGG (Figure 5A). However, given that cation coordination involves multiple bases (eight per cation), it is not possible to pinpoint a precise cation location on G-quadruplexes using EPD.

**Figure 5.**
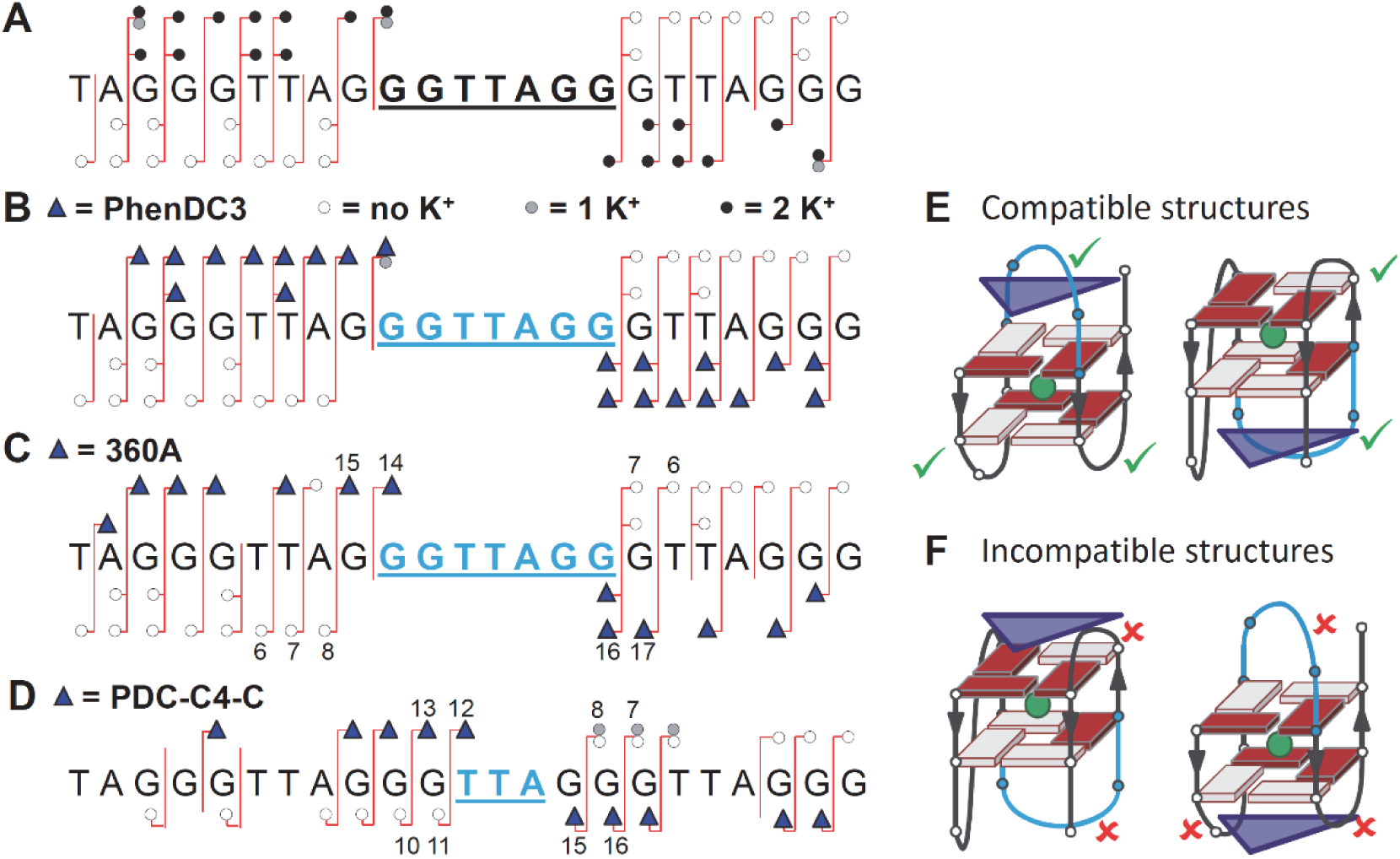
MS/MS of the complexes with 23TAG. A) EPD analysis of the MK_2_^6-^ complex. EPD analysis of the MKL^6-^ complexes with PhenDC3 (B) and 360A (C). D) CID analysis of the MKL^6-^ complex with cross-linked PDC-C4-C. In B—D, the violet triangles represent the ligand. On the right are represented the 2-quartet antiparallel complex topologies that are compatible (E) or incompatible (F) with ligand location close to the central loop.

EPD analysis was then carried out on the complexes with the noncovalently bound ligands 360A and PhenDC3. The complexes MKL fragment very similarly to the MK_2_ complexes (Figure 5 B-C for M = 23TAG; Supporting Figure S7 for 22GT): both L and K^+^ were detected on fragments including the central G_10_GTTAGG_16_ sequence, and never on fragments not including that sequence. This holds both for 360A and PhenDC3. Ligand binding close to the central loop is compatible with several complex topologies (Figure 5E) and allows to exclude others (Figure 5F). Thus, although the protection of a 7-nt stretch from fragmentation due to the complex formation precludes a precise location of the ligand, EPD provides useful information to outline which binding topologies are plausible or not for each complex.

To validate the central loop ligand location obtained by EPD of the noncovalent complex, we compared the results with the alkylating approach. To this aim, chlorambucil derivatives of PhenDC3 and 360A were synthesized (Figure 1D for PDC-C4-C and supporting information for more detail). Chlorambucil has two electrophilic sites and may induce both mono- and bis-alkylation on DNA, reacting mainly with the N7 of the guanines.^51, 52^ The ligands were allowed to react with their target (G-quadruplex pre-folded in 1 mM KCl) at room temperature for 24 hours. MS analysis highlights one of the main caveats of the cross-linking approach: the chlorambucil derivatives do not bind their target DNA with the same affinity as the original noncovalent ligands: PhenDC3-C4-C binds >100 times more poorly than PhenDC3, and PDC-C4-C binds with 10-fold lower affinity than 360A (Supporting Figure S8). PDC-C4-C binds exclusively by producing DNA mono-alkylation (loss of one HCl molecule) and ejection of one potassium ion (the MLK complex predominates, see Supporting Figure S9). So, despite the lower affinity of PDC-C4-C to G4, we assume that its binding mode is similar to that of 360A.

EPD analysis was not feasible on the MLK complex of PDC-C4-C: as the ligand binds by mono-alkylation, one chlorine group remains in the complex; as a result, the 5-● ions fragment exclusively by chlorine radical loss. CID was therefore performed on the 5- and 6- ions with 22GT and 23TAG (supporting Figure S10). The detected fragments are summarized in Figure 5D for the MLK^6-^ complex with 23TAG. The shortest 5’-terminal fragment on which L is detected is *a*_15_-Base, and given that the lost base is G15, it means that L is located somewhere on the first 14 bases. Similarly, the shortest 3’-fragment containing L is *w*_12_^z-^ (complementary to *a*_11_^z-^). This allows to bracket the cross-linking site on the central TTA loop.

Given the preferred reactivity of chlorambucil to guanines, binding to the loop sounds unusual. In a previous report, Balasubramanian group reported alkylation of a close analogue (pyridostatin-chlorambucil conjugate)^53^ to telomeric DNA, and detected after nuclease digestion that the ligand was covalently bound to both adenines and guanines. To confirm the MS/MS results, we performed gel sequencing analysis on PDC-C4-C adducts on the sequence 22AG (dAGGG(TTAGGG)_3_), the same as used by the Balasubramanian group. Alkaline treatment of the PDC-C4-C adducts revealed exclusive alkylation on thymine nucleobase localized within the central loop and on the two proximal guanines (Supporting Figure S11). All these results support the conclusions we had drawn from top-down EPD on the noncovalent underivatized complexes, namely that ligands of the PDC family are located close to the central loop of 4-repeat telomeric sequences.

## CONCLUSIONS

In summary, we have demonstrated how electron photodetachment dissociation (EPD), a top-down mass spectrometry sequencing approach, can be used to determine noncovalent binding sites on nucleic acids. Because the method does not require ligand cross-linking, tedious ligand derivatization procedures and risks of modifying the ligand behavior are alleviated. Localization is also possible on binders that are impossible to derivatize (e.g., single cations). Importantly, with mass spectrometry, top-down analysis can be carried on each complex stoichiometry individually, even though a mixture of stoichiometries may exist in solution. Enabling EPD on higher-resolution mass spectrometers and automating the fragment annotation will increase the applicability of the technique to nucleic acid complex characterization, in a similar way as ECD or ETD have now become routine techniques to localize post-translational modification on proteins.

## REFERENCES

1. S. Burge, G. N. Parkinson, P. Hazel, A. K. Todd and S. Neidle, Nucleic Acids Res., 2006, 34, 5402–5415.

2. A. I. Karsisiotis, C. O’Kane and M. Webba da Silva, Methods, 2013, 64, 28–35.

3. S. A. Dvorkin, A. I. Karsisiotis and M. Webba da Silva, Sci. Adv., 2018, 4, eaat3007.

4. J. Dai, M. Carver and D. Yang, Biochimie, 2008, 90, 1172–1183.

5. J. B. Chaires, FEBS J., 2010, 277, 1098–1106.

6. A. T. Phan, FEBS J., 2010, 277, 1107–1117.

7. R. D. Gray, J. O. Trent and J. B. Chaires, J. Mol. Biol., 2014, 426, 1629–1650.

8. R. D. Gray and J. B. Chaires, Biophys. Chem., 2011, 159, 205–209.

9. M. C. Miller, R. Buscaglia, J. B. Chaires, A. N. Lane and J. O. Trent, J. Am. Chem. Soc., 2010, 132, 17105–17107.

10. D. Miyoshi, K. Nakamura, H. Tateishi-Karimata, T. Ohmichi and N. Sugimoto, J. Am. Chem. Soc., 2009, 131, 3522–3531.

11. K. W. Lim, S. Amrane, S. Bouaziz, W. Xu, Y. Mu, D. J. Patel, K. N. Luu and A. T. Phan, J. Am. Chem. Soc., 2009, 131, 4301–4309.

12. M. Lenarcic Zivkovic, J. Rozman and J. Plavec, Angew. Chem. Int. Ed., 2018, 57, 15395–15399.

13. S. Balasubramanian, L. H. Hurley and S. Neidle, Nat. Rev. Drug Discov., 2011, 10, 261–275.

14. A. Cammas and S. Millevoi, Nucleic Acids Res., 2016, 45, 1584–1595.

15. J. F. Riou, Curr. Med. Chem., 2004, 4, 439–443.

16. S. Neidle, J. Med. Chem., 2016, 59, 5987–6011.

17. E. Ruggiero and S. N. Richter, Nucleic Acids Res., 2018, 46, 3270–3283.

18. S. Müller and R. Rodriguez, Expert Rev. Clin. Pharmacol., 2014, 7, 663–679.

19. A. R. Duarte, E. Cadoni, A. S. Ressurreicao, R. Moreira and A. Paulo, ChemMedChem, 2018, 13, 869–893.

20. P. Chilka, N. Desai and B. Datta, Molecules, 2019, 24, 752.

21. A. Marchand, A. Granzhan, K. Iida, Y. Tsushima, Y. Ma, K. Nagasawa, M. P. Teulade-Fichou and V. Gabelica, J. Am. Chem. Soc., 2015, 137, 750–756.

22. M. J. Lecours, A. Marchand, A. Anwar, C. Guetta, W. S. Hopkins and V. Gabelica, Biochim. Biophys. Acta, 2017, 1861, 1353–1361.

23. A. Marchand, F. Rosu, R. Zenobi and V. Gabelica, J. Am. Chem. Soc., 2018, 140, 12553–12565.

24. A. T. Phan, V. Kuryavyi, K. N. Luu and D. J. Patel, Nucleic Acids Res., 2007, 35, 6517–6525.

25. K. Breuker, M. Jin, X. Han, H. Jiang and F. W. McLafferty, J. Am. Soc. Mass. Spectrom., 2008, 19, 1045–1053.

26. K. B. Turner, N. A. Hagan, A. S. Kohlway and D. Fabris, J. Am. Soc. Mass Spectrom., 2006, 17, 1401–1411.

27. E. M. Schneeberger and K. Breuker, Angew. Chem. Int. Ed., 2017, 56, 1254–1258.

28. J. Wu and S. A. McLuckey, Int. J. Mass Spectrom., 2004, 237, 197–241.

29. J. P. Williams, J. M. Brown, I. Campuzano and P. J. Sadler, Chem. Commun., 2010, 46, 5458–5460.

30. Y. Xie, J. Zhang, S. Yin and J. A. Loo, J. Am. Chem. Soc., 2006, 128, 14432–14433.

31. S. Yin and J. A. Loo, Int. J. Mass Spectrom., 2011, 300, 118–122.

32. E. Boeri Erba, Proteomics, 2014, 14, 1259–1270.

33. K. F. Haselmann, T. J. D. Jorgensen, B. A. Budnik, F. Jensen and R. A. Zubarev, Rapid. Commun. Mass Spectrom., 2002, 16 2260–2265.

34. J. Yang and K. Hakansson, J. Am. Soc. Mass Spectrom, 2006, 17, 1369–1375.

35. V. Gabelica, F. Rosu, T. Tabarin, C. Kinet, R. Antoine, M. Broyer, E. De Pauw and P. Dugourd, J. Am. Chem. Soc., 2007, 129, 4706–4713.

36. S. Daly, M. Porrini, F. Rosu and V. Gabelica, Faraday Discuss., 2019, 10.1039/C1038FD00207J.

37. V. Gabelica, T. Tabarin, R. Antoine, F. Rosu, I. Compagnon, M. Broyer, E. De Pauw and P. Dugourd, Anal. Chem., 2006, 78, 6564–6572.

38. V. Larraillet, R. Antoine, P. Dugourd and J. Lemoine, Anal. Chem., 2009, 81, 8410–8416.

39. R. Antoine, J. Lemoine and P. Dugourd, Mass Spectrom. Rev., 2014, 33, 501–522.

40. M. J. Cavaluzzi and P. N. Borer, Nucleic Acids Res., 2004, 32, e13.

41. A. Marchand and V. Gabelica, J. Am. Soc. Mass Spectrom., 2014, 25, 1146–1154.

42. A. Marchand and V. Gabelica, Nucleic Acids Res., 2016, 44, 10999–11012.

43. A. T. Iavarone, J. C. Jurchen and E. R. Williams, Anal. Chem., 2001, 73, 1455–1460.

44. H. J. Sterling, M. P. Daly, G. K. Feld, K. L. Thoren, A. F. Kintzer, B. A. Krantz and E. R. Williams, J. Am. Soc. Mass Spectrom., 2010, 21, 1762–1774.

45. B. Brahim, S. Alves, R. B. Cole and J. C. Tabet, J Am Soc Mass Spectrom, 2013, 24, 1988–1996.

46. N. Xu, K. Chingin and H. Chen, J. Mass Spectrom., 2014, 49, 103–107.

47. R. R. Ogorzalek Loo, R. Lakshmanan and J. A. Loo, J. Am. Soc. Mass Spectrom., 2014, 25, 1675–1693.

48. S. Ickert, J. Hofmann, J. Riedel, S. Beck, K. Pagel and M. W. Linscheid, Eur. J. Mass Spectrom., 2018, 24, 225–230.

49. M. Porrini, F. Rosu, C. Rabin, L. Darre, H. Gomez, M. Orozco and V. Gabelica, ACS Cent. Sci., 2017, 3, 454–461.

50. N. Khristenko, J. Amato, S. Livet, B. Pagano, A. Randazzo and V. Gabelica, J. Am. Soc. Mass Spectrom, 2019, DOI: 10.1007/s13361-019-02152-3.

51. W. B. Mattes, J. A. Hartley and K. W. Kohn, Nucleic Acids Res., 1986, 14, 2971–2987.

52. J. A. Hartley, J. P. Bingham and R. L. Souhami, Nucleic Acids Res., 1992, 20, 3175–3178.

53. M. Di Antonio, K. I. McLuckie and S. Balasubramanian, J. Am. Chem. Soc., 2014, 136, 5860–5863.

