## supporting information for "Probing Ligand and Cation Binding Sites in G-Quadruplex Nucleic Acids by Mass Spectrometry and Electron Photodetachment Dissociation Sequencing"

#### Supporting Information - Table of Contents

Additional experimental details (synthesis of the ligands, circular dichroism and gel analysis).

Figure S1: Annotation of DNA-cation stoichiometries observed in full scan MS spectra (without ligand).

Figure S2: DNA-cation-ligand stoichiometries in full scan MS spectra with ligands 360A and PhenDC3.

Figure S3: Comparison of EPD on $M^{5-}$ and $M^{6-}$ (DNA without cation or ligand).

Figure S4: Circular dichroism spectra of the complexes in the absence and presence of sulfolane.

Figure S5: Comparisons between EPD and CID fragmentation.

Figure S6: EPD of 23TAG with zero, one and two potassium ions ($M^{6-}$, $MK^{6-}$ and $MK_2^{6-}$).

Figure S7: EPD of 22GT with ligands PhenDC3 and 360A.

Figure S8: ESI-MS characterization of the binding of the chlorambucil derivatives to 23TAG (affinity, loss of single chlorine, ejection of potassium).

Figure S9: Isotopic pattern analysis proving that PDC-C4-C binds with ejection of one potassium from the DNA and loss of one chlorine atom from the cross-linked ligand.

Figure S10: EPD and CID on the complexes of 23TAG and 22GT with PDC-C4-C.

Figure S11: Analysis of cross-linking by gel electrophoresis.

### Experimental details:

**Ligands synthesis and characterization.** $^{1}H$ and $^{13}C$ spectra were recorded at 25°C on a Bruker Avance 300 using TMS as internal standard. Deuterated $CDCl_3$ and $[D_6]DMSO$ were purchased from SDS. The following abbreviations are used: singlet (s), doublet (d), triplet (t) and multiplet (m). Low resolution mass spectrometry (ESI-MS) was recorded on a micromass ZQ 2000 (waters). High resolution mass spectrometry (ESI-MS) and $^{13}C$ NMR were provided by the I.C.S.N. (Institut de Chimie des Substances Naturelles, Gif-sur-Yvette). TLC analysis was carried out on silica gel (Merck 60F-254) with visualization at 254 and 366 nm. Melting points were taken on a Kofler melting point apparatus and are uncorrected. Preparative flash chromatography was carried out with Merck silica gel (Si 60A, 35-70 µm) using Dichloromethane/Methanol mixtures unless otherwise state. Reagents and chemicals were purchased from Sigma-aldrich, Acros or Alfa-aesar unless otherwise stated: chelidamic acid, 3-aminoquinoline, 1,4-butanediamine, and chlorambucil are commercially available. Solvents were purchased from SDS.

**Synthesis of PDC-C4-C.**

To achieve the synthesis of the alkylating derivative PDC-C4-C, we exploit an easy procedure starting from the precursor **1**, obtained in four steps following a previous described synthetic protocol.[1,]

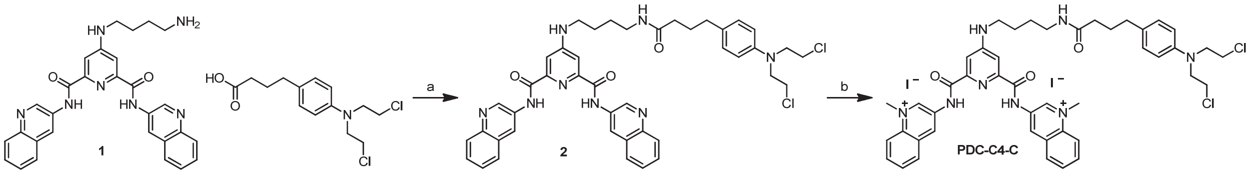


**Scheme S1:** Synthesis of the intermediate compound **2**, and final compound PDC-C4-C: a) 1.0 eq. **1**, 2.0 eq. chlorambucil, 2.0 eq. HBTU, 5.0 eq. DIEA, in DMF at r.t. 24 h, 93%; b) $CH_3I$ (1.50 mL), DMF (1.0 ml), 40 °C, 24 h.

**4-((4-(4-(4-(bis(2-chloroethyl)amino)phenyl)butanamido)butyl)amino)-N2,N6-di(quinolin-3-yl)pyridine-2,6-dicarboxamide (2)**

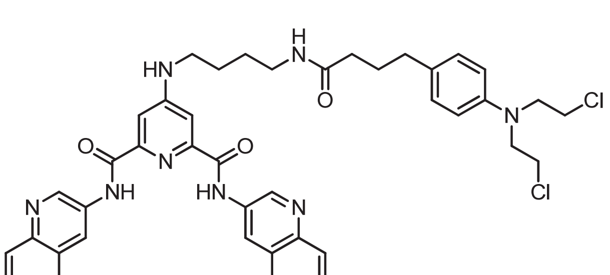

In a 20 mL round-bottomed flask was added 4-((4-aminobutyl)amino)-N2,N6-di(quinolin-3-yl)pyridine-2,6-dicarboxamide (**1**) (100 mg, 0.198 mmol), chlorambucil 120 mg, 0.396 mmol), and DIEA (0.164 ml, 38.79 mmol) in DMF (3.0 ml) to give a colorless solution. Within stirring HBTU (150 mg, 0.396 mmol) was added. The solution was stirred for 24 h at room temperature. The reaction was then diluted with water and extracted with ethyl acetate (3X50 mL). The combined organic extract was dried over $MgSO_4$. The crude product was added to a silica gel column and eluted with acetate/hexanes 1:1. Purification afforded compound **2** as a white solid (146mg, 93 % yield). Dec. **>** 124°C; **$^{1}$H NMR** (300 MHz, $CDCl_3$): $\delta$= 10.40 (s, 2H), 9.10 (s, 2H), 8.56 (s, 2H), 7.90 (d, $^{3}J$(H,H)= 6 Hz, 2H), 7.53-7.32 (m, 8H), 6.90 (d, $^{3}J$(H,H)= 9 Hz, 2H), 6.48 (d, $^{3}J$(H,H)= 9 Hz, 2H), 6.29

1 Renaud de la Faverie, A., Hamon, F., Di Primo, C., Largy, E., Dausse, E., Delaurière, L., Landras-Guetta, C., Toulmé, J.-J., Teulade-Fichou, M.-P., Mergny, J.-L. *Biochimie* **2011**, *93*, 1357-1367.

(ds, 2H), 3.61-3.54 (m, 8 H), 3.20 (bs, 2H), 3.02 (bs, 2H), 2.40 (t, J = 7.5 Hz, 2H), 2.09-2.07 (m, 2H), 1.84-1.78 (m, 2H), 1.51 (bs, 4H) ppm; **$^{13}$C NMR** (75 MHz, $CDCl_3$): $\delta$= ppm; **LRMS** (ESI-MS): m/z=791.3 $[M+H]^+$, 396.2 [M+2H]2+, 815.3 $[M+Na]^+$. **HRMS** (ESI-MS): 791.3003 (calculated: 791,2992 $C_{43}H_{44}Cl_2N_8O_3^+$).

**3,3'-((4-((4-(4-(4-(bis(2-chloroethyl)amino)phenyl)butanamido)butyl)amino)pyridine-2,6-dicarbonyl)bis(azanediyl))bis(1-methylquinolin-1-ium) iodide (PDC-C4-C)**

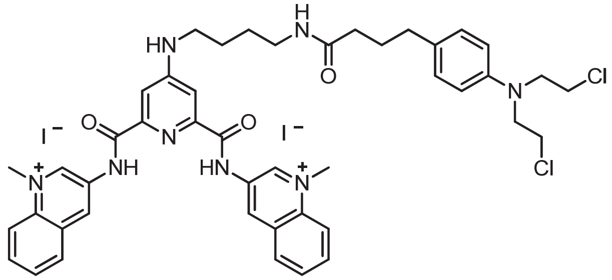

In a 10 mL round-bottomed flask, compound **2** (52.mg, 0.0657 mmol) was dissolved in DMF (1.00 mL) to give a colorless solution. $CH_3I$ (1.5 mL, 24.09 mmol) was added. The reaction was stirred overnight at 40 °C. The reaction mixture was let to cool down and poured in ethyl ether. The yellow solid was filtered and washed several times with tertbutylmethyl ether. The product was left to dry. The product PDC-C4-C has been obtained as yellow solid (47.5 mg, 67% yield). Dec. > 149°C; **$^1$H NMR** (300 MHz, $[D_6]$DMSO): $\delta$= 11.74 (s, 2H), 10.14 (s, 2H), 9.65 (s, 2H), 8.56 (d, $^3J$(H,H)= 9 Hz, 2H), 8.24 (t, $^3J$(H,H)= 7.5 Hz, 2H), 8.08 (t, $^3J$(H,H)= 7.5 Hz, 2H), 7.84 (t, $^3J$(H,H)= 6 Hz, 1H), 7.72 (t, $^3J$(H,H)= 6 Hz, 1H), 7.60 (bs, 2H), 7.00 (d, $^3J$(H,H)= 9 Hz, 2H), 6.64 (d, $^3J$(H,H)= 9 Hz, 2H), 4.78 (s, 6H), 3.68 (s, 8 H), 3.2 (m, 2H), 3.11 (m, 2H), 2.43 (t, J = 6 Hz, 2H), 2.07 (t, t, J = 6 Hz, 2H), 1.78-1.54 (m, 6H) ppm; **$^{13}$C NMR** (75 MHz, $[D_6]$DMSO): $\delta$= 163.1, 156.2, 146.0, 144.6, 132.0, 128.6, 128.2, 127.8, 127.7, 127.0, 124.0, 42.0, 41.2, 32.3, 29.9, 28.9, 28.8, 28.1, 26.5, 26.3ppm; **LRMS** (ESI-MS): m/z= 947.74 $[M-I]^+$, 410.47 $[M+2H]^+$. **HRMS** (ESI-MS): 947.2427 (calculated: 947.2428 $C_{45}H_{50}Cl_2IN_8O_3^+$).

**Synthesis of PhenDC3-C4-C**

To achieve the synthesis of the alkylating derivative PhenDC3-C4-C, we exploit an easy procedure starting from the precursor **3**, obtained in eight steps following a previous described synthetic protocol.[2,]

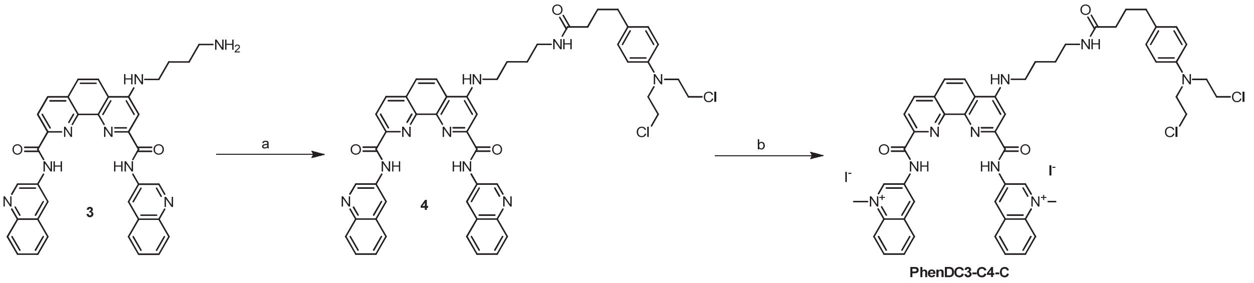


**Scheme S2:** Synthesis of the final product PhenDC3-C4-C: a) chlorambucil (2 eq), DIEA (5 eq), and HBTU (2 eq) in DMF rt for 20 h; b) iodomethane in DMF 40°C (41%).

2 Lefebvre, J., Guetta, C., Poyer, F., Mahuteau-Betzer, F., Teulade-Fichou, M. P., *Angew. Chem. Int. Ed.* **2017**, *56*, 11365-11369.

**4-((4-(4-(4-(bis(2-chloroethyl)amino)phenyl)butanamido)butyl)amino)-N2,N9-di(quinolin-3-yl)-1,10-phenanthroline-2,9-dicarboxamide (4)**

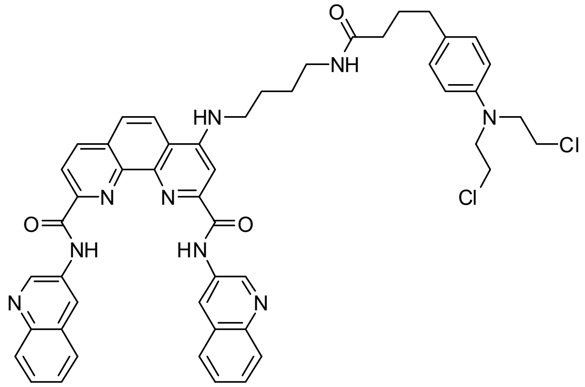

A 10 mL flask was charged with a magnetic spin bar, compound **3** (67 mg, 0.110 mmol), chlorambucil (67.19 mg, 0.220 mmol), DMF( 1.67 mL), and diisopropylethylamine (0.91 mL, 0.550 mmol). Under stirring HBTU (83.77 mg, 0.22 mmol) was added and the reaction was stirred for 20 h at room temperature. The solvent was removed under reduced pressure and the residue purificied by $SiO_2$ chromatography DCM-MeOH 100/00 to 96/5. Product **4** was obtained as a yellow powder (29 mg, 29%). Dec> 160°C°; **$^1$H NMR** (300 MHz, [D6]DMSO): δ= 11.84 (m,2H), 9.67(bs,2H), 9.14 (m,2H), 8.78(m,1H), 8.61-8.55(m,2H), 8.09(m, 4H), 7.87-7.66(m,7H), 6.98(d,9Hz,2H), 6.59(d,6Hz,2H), 3.65(s,8H), 3.51(m, 3H), 2.09(m,3H), 1.77-1.62(m, 6H), 0.85(m,2H) ppm; **$^{13}$C NMR** (75 MHz, [D6]DMSO): δ= ppm; **LRMS** (ESI-MS): m/z= 892$[M+H]^+$. **HRMS** (ESI-MS): (calculated:$C_{50}H_{47}Cl_2N_9O_3$).

**3,3'-((4-((4-(4-(4-(bis(2-chloroethyl)amino)phenyl)butanamido)butyl)amino)-1,10-phenanthroline-2,9-dicarbonyl)bis(azanediyl))bis(1-methylquinolin-1-ium) iodide (PhenDC3-C4-C)**

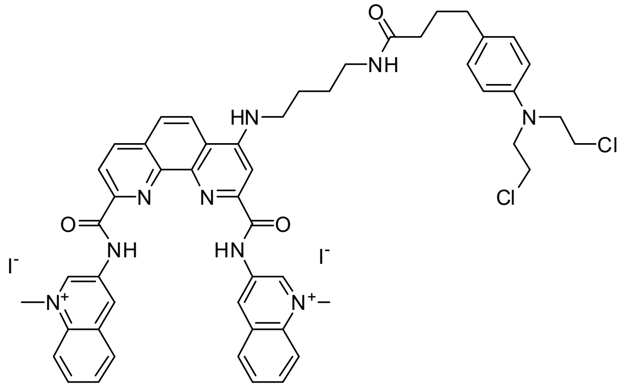

To the solution of compound **4** (21.38 mg, 0.024 mmol) in dry DMF (0.69 mL) heated at 40°C was added dropwise an excess of methyl iodide ( 59.6 µL, 0.96 mmol).

The reaction mixture was stirred under argon atmosphere and in the dark for 16 h at 40°C. The reaction mixture was let to cool down and poured in ethyl ether. The obtained solid was filtered from diethyl ether and dried. The product was obtained as an orange powder which was dried under vacuum (28 mg, 99%). **Dec** > 90°C °C; **$^1$H NMR** (300 MHz, $[D_6]$DMSO): $\delta$= 12.14 (bs, 2H), 10.32 (s, 2H), 9.88 (s, 2H), 8.90 (m, 1H), 8.70-8.51 (m, 5H), 8.24-8.05 (m, 10H), 7.00 (d, $J$=9 Hz, 2H), 6.60 (m, 2H), 4.72 (s, 6H), 3.66 (s, 8H), 3.24-3.16 (m, 5H), 2.09 (m, 2H), 1.78-1.62 (m, 7H).ppm. **$^{13}$C NMR** (75 MHz, $[D_6]$DMSO): $\delta$= ppm. **LRMS** (ESI-MS): m/z= 460.7 $[M-2\ I\ /2]^{2+}$. **HRMS** (ESI-MS): 921.3608 (calculated: 921.3648 $C_{52}H_{53}Cl_2N_9O_3^{2+}$).

**Circular dichroism (CD) spectroscopy.** Circular dichroism experiments were run on a Jasco J815 spectrophotometer. The samples were prepared 36 hours in advance, including the addition of sulfolane or ligand if any. Shown spectra are the results of three accumulations using a quartz cell of 2 mm optical path length at 20 °C with a scan speed of 50 nm/min and 0.5 s integration time. The raw data were normalized to molar-circular dichroic absorption Δε based on the DNA concentration (C = 10 $x10^{-6}$ M): Δε = θ/(32980 x l x C), where θ is the ellipticity in milidegrees and l, the optical path length (l = 0.2 cm). The data were baseline subtracted using a 100 mM TMAA in pure water solution in the same cuvette as the experiment.

**Gel analysis.** The alkylation reactions were performed by using solutions of 20 µL in 2 mL centrifuge tubes. The tubes were placed into a hot plate at 37 °C for 16 h. 22AG d[A(GGGTTA)$_3$GGG] was synthesized and purified by Eurogentec, desalted on a Sephadex G25 column and stored at -20 °C as a 250 µM aqueous solution. The oligonucleotide was 5'-end-labeled using bacterial T4 polynucleotide kinase (purchased Pharmacia Biotech) and [$\gamma$-$^{32}$P]ATP (purchased Pharmacia Biotech). The labeling was performed according to standard procedures. Radiolabeled DNAs were purified by 20% denaturing gel electrophoresis.

**5'-End Labeling of Oligonucleotides.** The oligonucleotides were 5'-end-labeled using polynucleotide kinase (Pharmacia Biotech) and [γ-$^{32}$P]ATP (Pharmacia Biotech). The reaction products were purified via 20% denaturing gel electrophoresis and desalted on a Sephadex G25 column.

**Alkylation of 22AG d[A(GGGTTA)$_3$GGG] in the Presence of $K^+$.** The 5'-end-radiolabeled 22AG was mixed with 10 µM non-radiolabeled material in KCl for 5 min at 90 °C and allowed to reach room temperature in 2 h to induce the formation of quadruplex structure. It was then incubated with 25 µM PDC-C4-C (1:2.5 molar ratio) in a total volume of 40 µl. The reactions were carried out for 16 h at 37 °C. The alkylated oligonucleotides were separated by polyacrylamide denaturing gel electrophoresis and located by autoradiography. They migrated differently according to their masses and charges. The alkylation sites were determined from piperidine cleavage.

**Determination and quantification of the alkylating agent binding sites by piperidine treatment.** The gel migrations of the alkylated oligonucleotides depend on the type of adducts. An increasing number of bound alkylating agents slow down the migration, essentially because each of them brings two additional charges. However, the migration is accelerated when the adduct is forming loops on the structure. After their separation by gel electrophoresis, the different products of alkylation were eluted from the gel and precipitated with cold ethanol. The nonalkylated oligonucleotides were treated with dimethylsulfate (DMS)/piperidine (probe of the free $N_7$ of the guanine base) in Maxam-Gilbert sequencing conditions: the oligonucleotides were dissolved in 19 µL of water and incubated with 1 µL of DMS for 1 min 30 sec at 37 °C. The alkylation products and the nonalkylated oligonucleotides, after DMS/piperidine treatments and precipitation, were incubated with 50 µL of 1 M piperidine aqueous solution at 90 °C for 25 min, which induces the cleavage at the alkylation sites with partial regeneration of the starting 22AG. After evaporation of the piperidine, the samples were washed four times with water and migrated by 20% denaturing gel electrophoresis. The gels were then analyzed, and the cleavage sites identified, by a Molecular Dynamics Phosphorimager with Imagequant software for data processing.

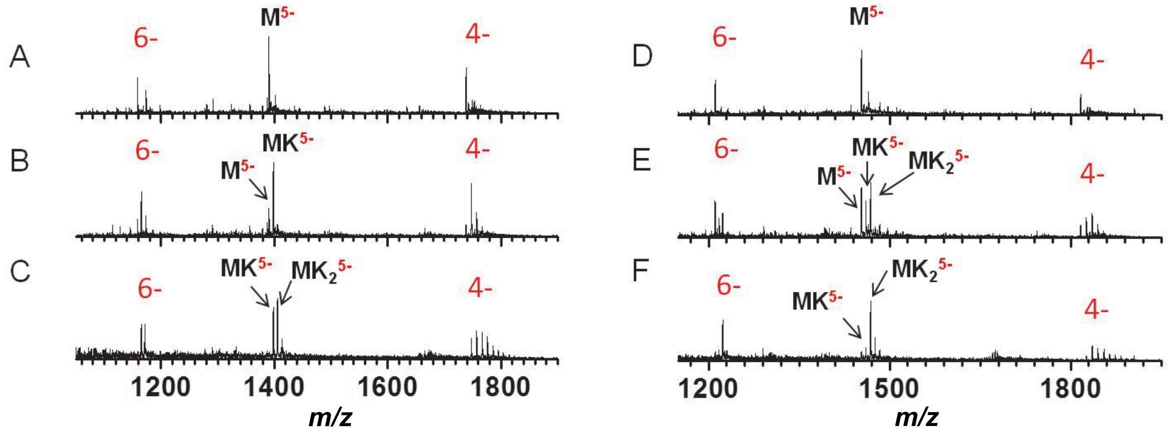


**Figure S1:** ESI-MS spectra of 10 µM 22GT/23TAG in 0 to 1 mM KCl/100 mM TMAA. A/D) 0 mM KCl, B/E) 0.2 mM KCl, and C/F) 1 mM KCl. A to C) for the sequence 22GT, and D to F) for the sequence 23TAG.

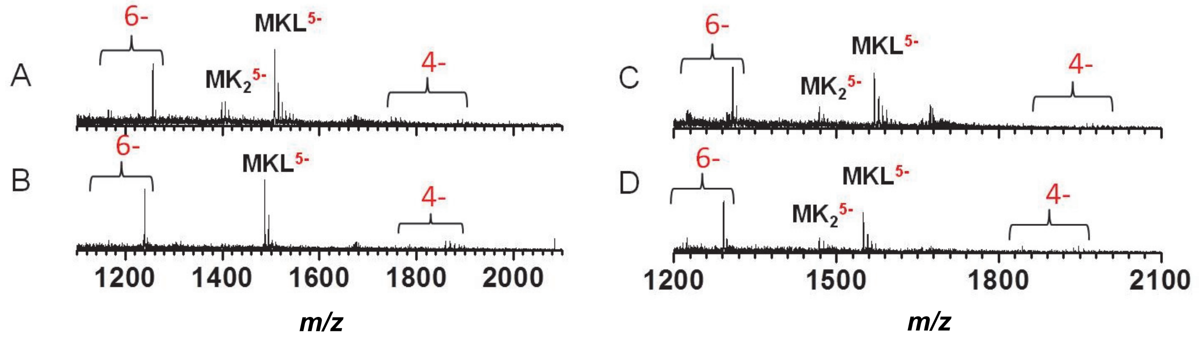


**Figure S2:** ESI-MS spectra of 10 µM 22GT/23TAG with equimolar concentration of ligands, PhenDC3 and 360A. All spectra were recorded in 1.0 mM KCl. A) sequence 22GT + ligand PhenDC3; B) sequence 22GT + ligand 360A; C) sequence 23TAG + ligand PhenDC3; D) sequence 23TAG + ligand 360A.

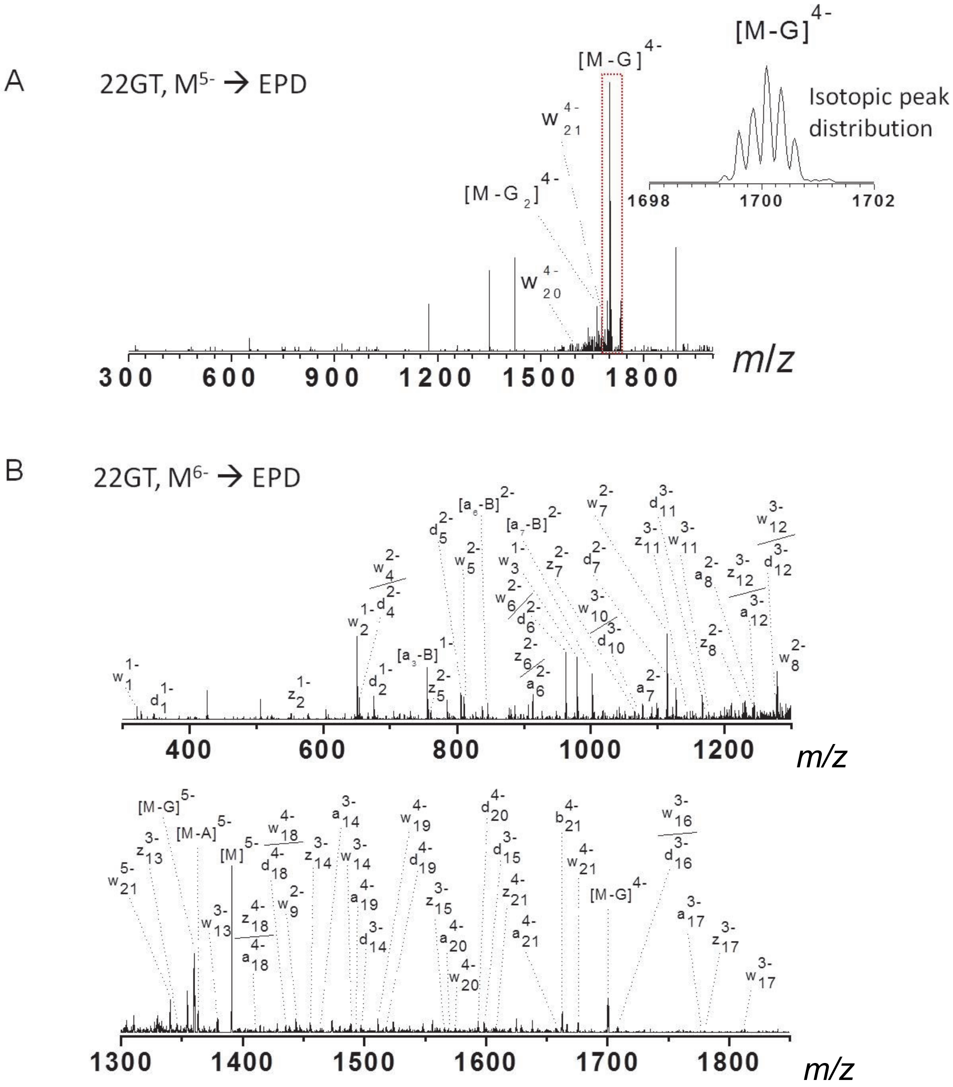


**Figure S3:** EPD on 5-(A) and 6-(B) for 22GT. Fragments are assigned manually based on their isotopic peak distribution as shown the inset of panel A.

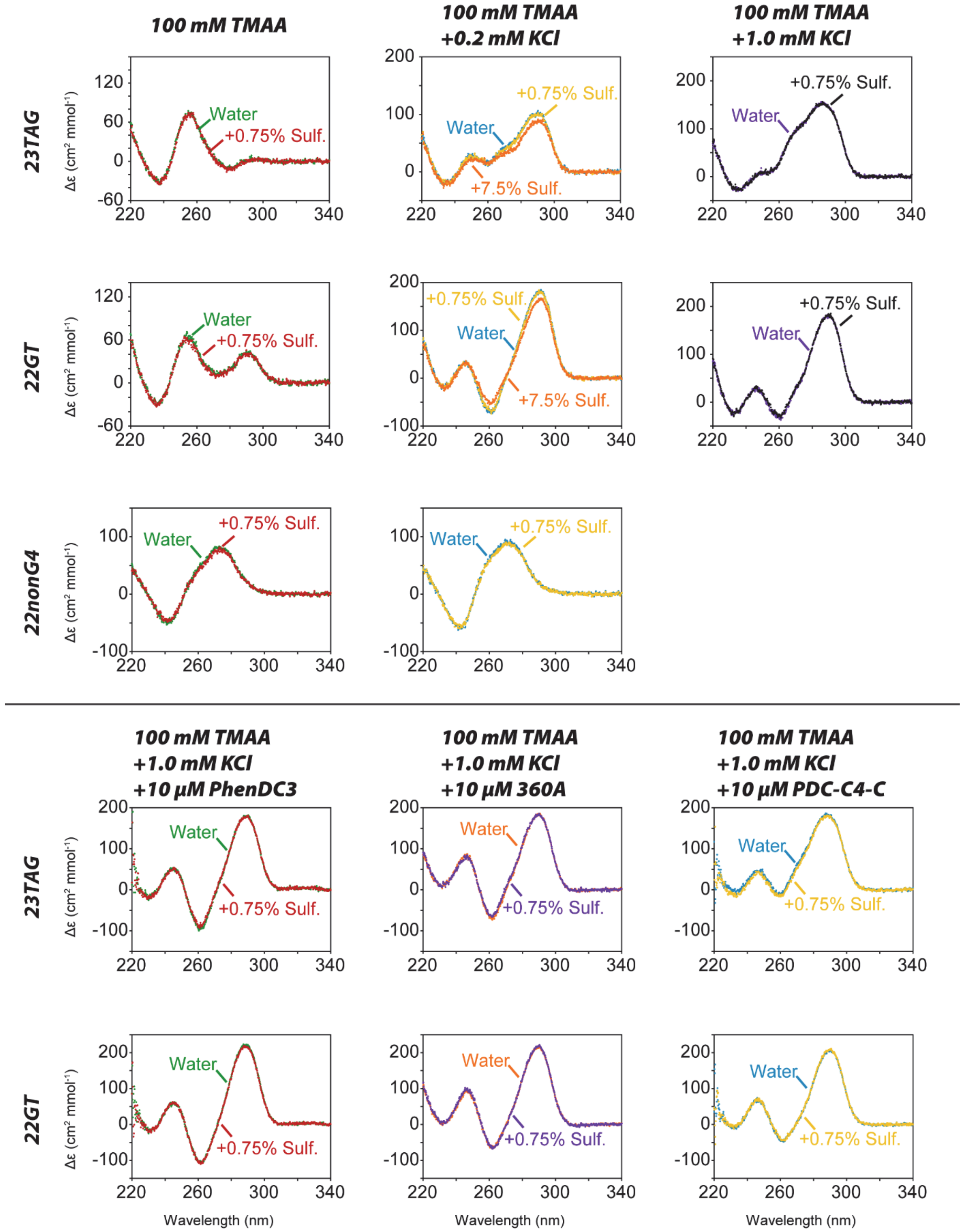


**Figure S4:** Circular dichroism spectra of the complexes in the absence and presence of sulfolane, showing that sulfolane does not alter the solution topology.

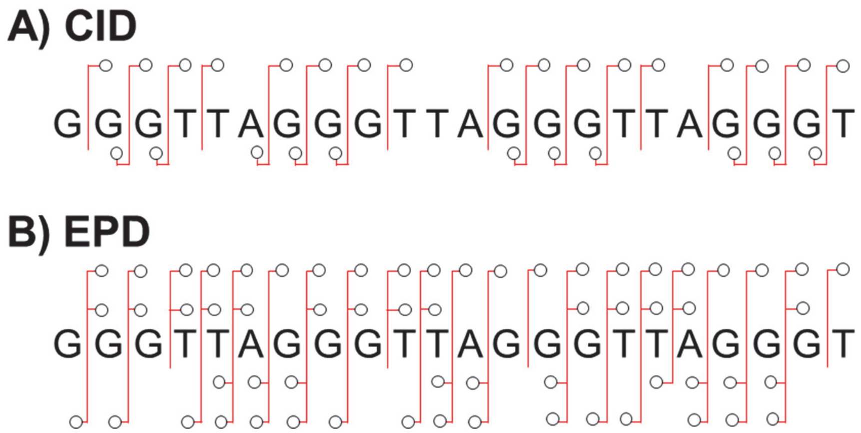


**Figure S5:** Comparison of the sequence fragmentation patterns of sequence $[22GT]^{6-}$ with (A) CID and (B) EPD.

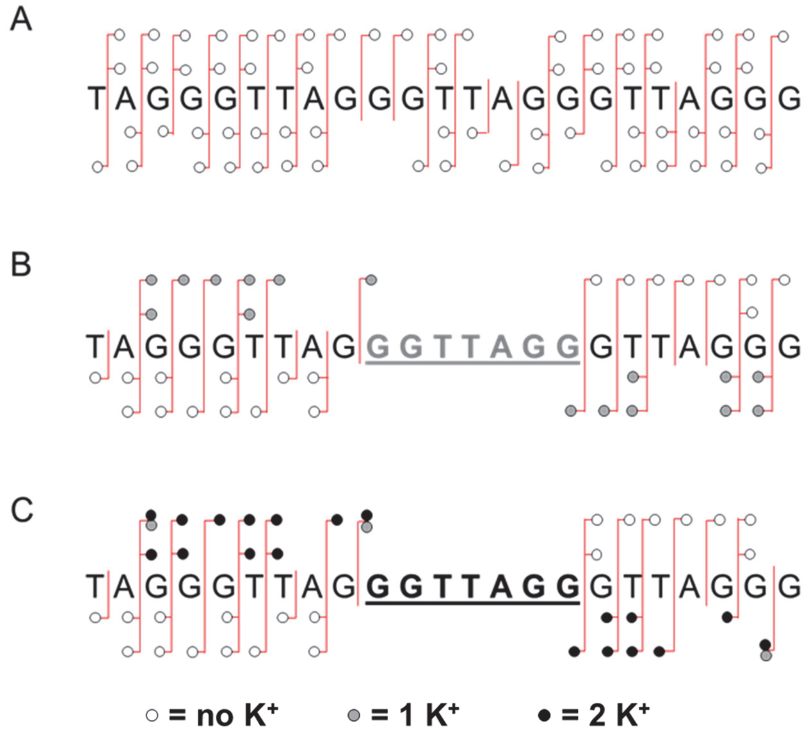


**Figure S6:** EPD of 23TAG with zero (A), one (B) and two potassium (C) ions ($M^{6-}$, $MK^{6-}$ and $MK_2^{6-}$, respectively).

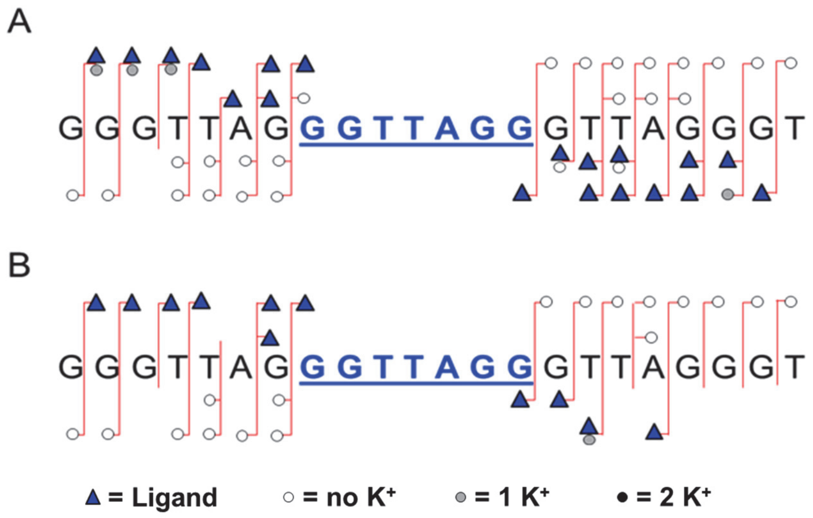


**Figure S7:** EPD sequencing of 22GT in its $MLK^{6-}$ complexes with ligands PhenDC3 (A) and 360A (B).

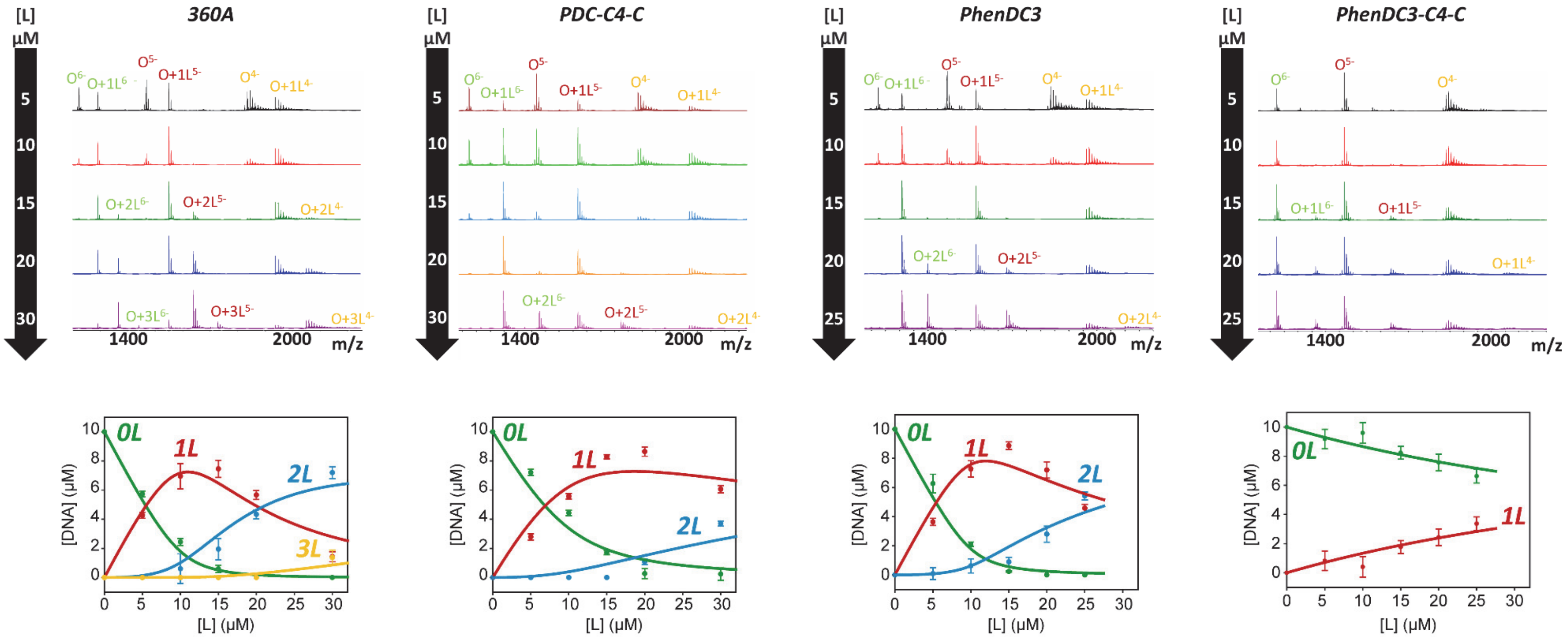


$K_{D1}$ = 0.19 ± 0.08 µM
$K_{D2}$ = 5.3 ± 0.9 µM

$K_{D1}$ = 1.6 ± 0.5 µM
$K_{D2}$ = 47 ± 13 µM

$K_{D1}$ = 0.27 ± 0.13 µM
$K_{D2}$ = 14 ± 3 µM

$K_{D1}$ = 56 ± 6 µM

**Figure S8:** Mass spectrometry titrations of 10 µM 23TAG with 360A, PDC-C4-C, PhenDC3 and PhenDC3-C4-C. The increasing ligand concentration is indicated in the black arrow. Bottom panel, quantification and fitting using the equations indicated below. The error bars are from the quantification of the different charge states. Equilibrium dissociation binding constants for the binding of the two first ligands are decreased by a factor 10 when 360A is derivatized with chlorambucyl and almost no binding is observed for PhenDC3-C4-C.

The equilibrium dissociation binding constants are defined as follow and fitted using the DynaFit software (4.05.103 BioKin Ltd Wa-tertown, MA) (Kuzmic, P. *Anal. Biochem.* **1996**, *237* (2), 260.):

$$DNA + L \rightleftharpoons DNA \bullet 1L \qquad K_{D1} = \frac{[DNA][L]}{[DNA \bullet 1L]}$$

$$DNA \bullet 1L + L \rightleftharpoons DNA \bullet 2L \qquad K_{D2} = \frac{[DNA \bullet 1L][L]}{[DNA \bullet 2L]}$$

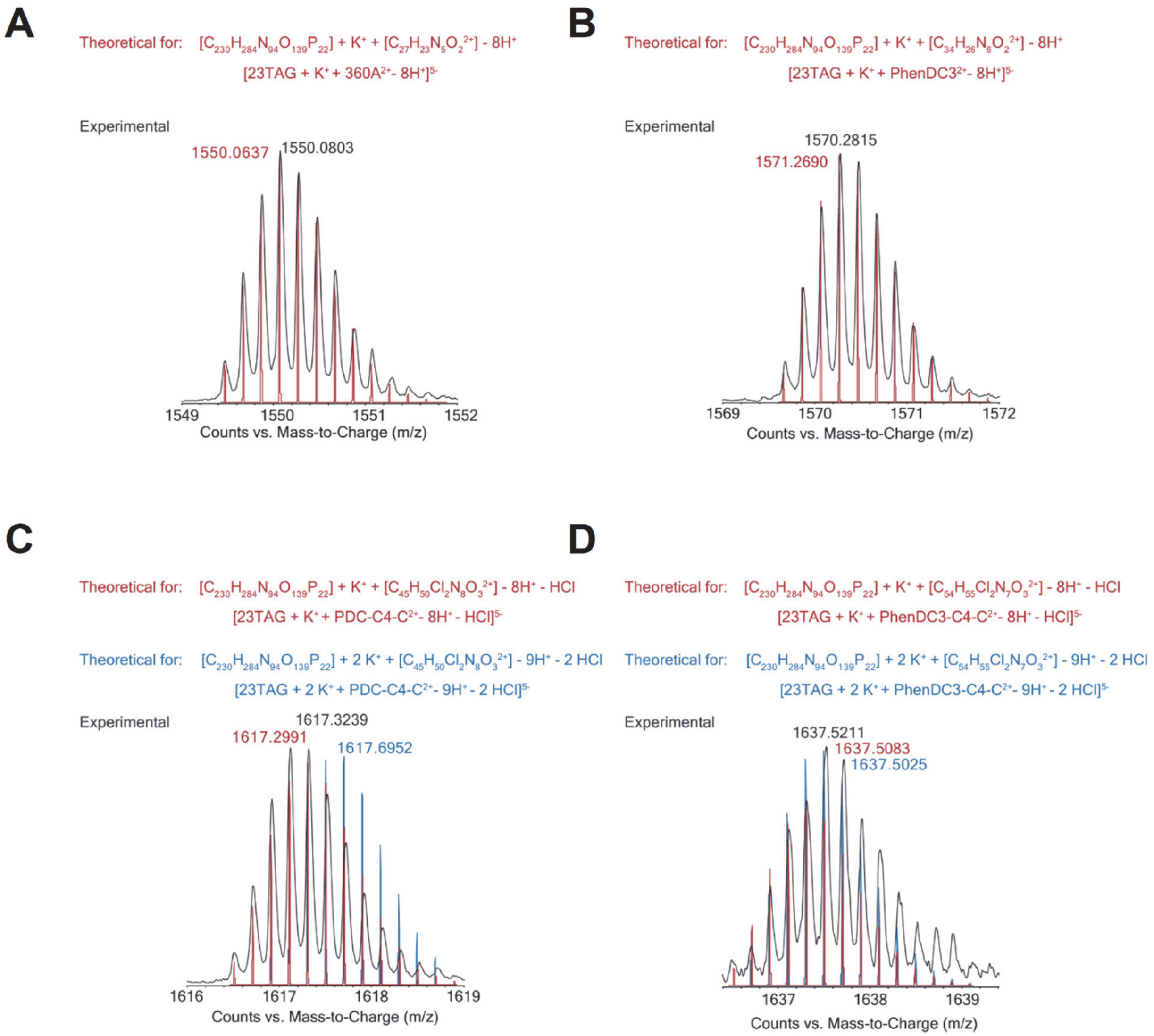


**Figure S9:** Zooms on the 1:1 (DNA:L) complexes. Identification of the major peak stoichiometry using theoretical isotopic distributions. 360A (A) and PhenDC3 (B) eject a potassium upon binding. PDC-C4-C (C) and PhenDC3-C4-C (D) also eject a potassium cation in addition with a HCl molecule, indicating the covalent bound between the DNA and the ligand. The difference between theoretical masses and experimental ones are about 10 ppm. For PDC-C4-C, the single loss of HCl is clear based on the isotopic distribution. For PhenDC3-C4-C, the discrimination is more difficult due to the weak abundance of the complex.

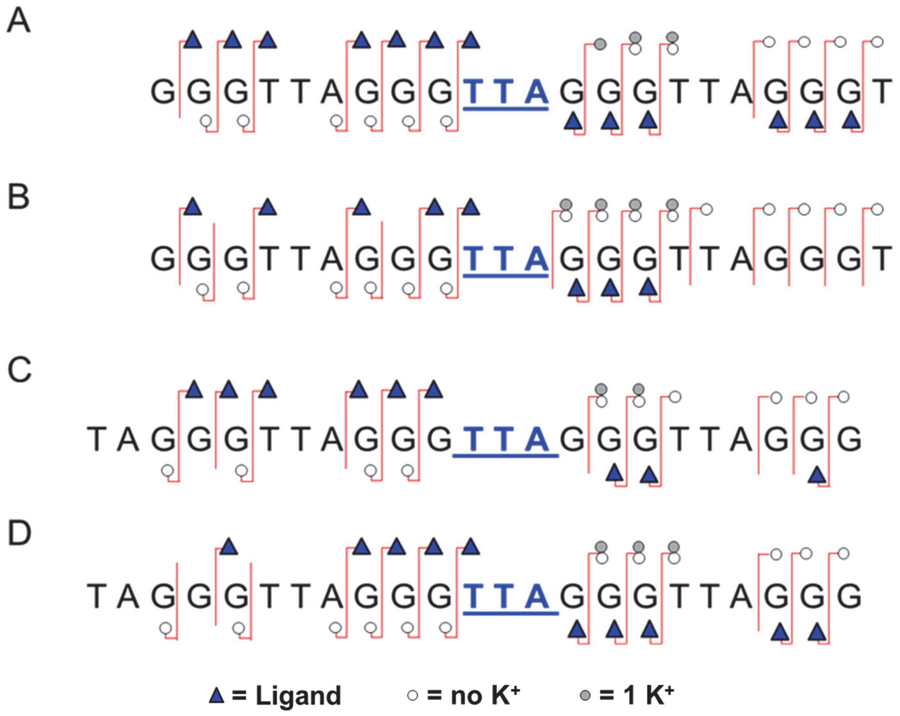


**Figure S10:** CID sequencing on the cross-linked complexes of PDC-C4-C with sequences (A-B) 22GT (charge state 5- in panel A and charge state 6- in panel B) and (C-D) 23TAG (charge state 5- in panel C and charge state 6- in panel D).

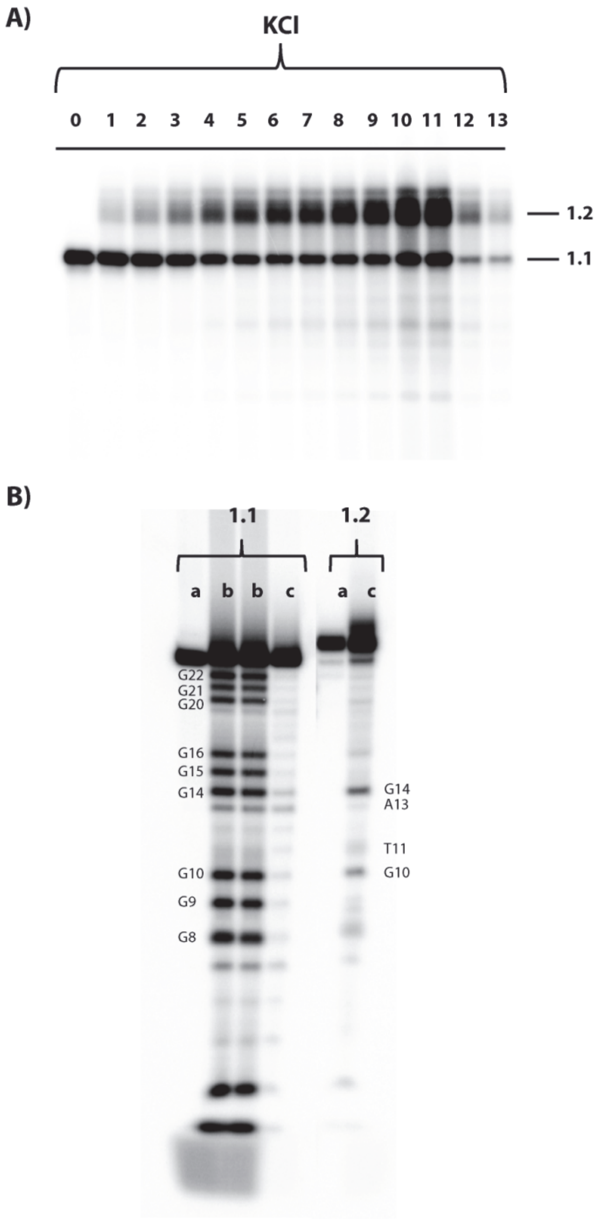


**Figure S11:** A) Denaturating gel electrophoresis (15% acrylamide) of the alkylation products of 22AG ($AG_3(T2AG_3)_3$ (10µM) by PDC-C4-C (25 µM) in the presence of KCl buffer (100 mM). Lane 0 is the control experiment that show the effect of no alkylation, (lanes 1-13) show the effects of alkylation at 37 °C after: 0.25, 0.5, 1, 2, 3, 4, 5, 6, 8, 10, 12, 15, 24 h. Bands have been numbered as function of migration as compared to control (band 1.1, no migration; band 1.2 retarded band). B) Analysis by sequencing gel electrophoresis (20% acrylamide) of the alkylation products of 22AG with PDC-C4-C isolated from the bands of the denaturing gel in $K^+$-rich buffer 100 mM from Figure S11A. Band 1.1, not alkylated 22AG and band 1.2 alkylated product; lane a: not treated sample; lane b: DMS+piperidine treatment, and it gives the reference scale for the migration of the guanines present in the 22AG sequence; lane c: piperidine treated sample. The numbered nucleobases correspond to the alkylated sites.
